# Phylogenetic and functional substrate specificity for endolithic microbial communities in hyper-arid environments

**DOI:** 10.1101/033340

**Authors:** Alexander Crits-Christoph, Courtney K. Robinson, Bing Ma, Jacques Ravel, Jacek Wierzchos, Carmen Ascaso, Octavio Artieda, Virginia Souza-Egipsy, M. Cristina Casero, Jocelyne DiRuggiero

## Abstract

Under extreme water deficit, endolithic (inside rock) microbial ecosystems are considered environmental refuges for life in cold and hot deserts, yet their diversity and functional adaptations remain vastly unexplored. The metagenomic analyses of the communities from two rock substrates, calcite and ignimbrite, revealed that they were dominated by *Cyanobacteria, Actinobacteria*, and *Chloroflexi*. The relative distribution of major phyla was significantly different between the two substrates and biodiversity estimates, from 16S rRNA gene sequences and from the metagenome data, all pointed to a higher taxonomic diversity in the calcite community. While both endolithic communities showed adaptations to extreme aridity and to the rock habitat, their functional capabilities revealed significant differences. ABC transporters and pathways for osmoregulation were more diverse in the calcite chasmoendolithic community. In contrast, the ignimbrite cryptoendolithic community was enriched in pathways for secondary metabolites, such as non-ribosomal peptides (NRPS) and polyketides (PKS). Assemblies of the metagenome data produced population genomes for the major phyla found in both communities and revealed a greater diversity of *Cyanobacteria* population genomes for the calcite substrate. Draft genomes of the dominant *Cyanobacteria* in each community were constructed with more than 93% estimated completeness. The two annotated proteomes shared 64% amino acid identity and a significantly higher number of genes involved in iron update, and NRPS gene clusters, were found in the draft genomes from the ignimbrite. Both the community-wide and genome-specific differences may be related to higher water availability and the colonization of large fissures and cracks in the calcite in contrast to a harsh competition for colonization space and nutrient resources in the narrow pores of the ignimbrite. Together, these results indicated that the habitable architecture of both lithic substrates-chasmoendolithic versus cryptoendolithic - might be an essential element in determining the colonization and the diversity of the microbial communities in endolithic substrates at the dry limit for life.

## Introduction

The rate of desertification across our planet is accelerating as the result of human activity and climate change. Ecosystems of arid and hyperarid environments are highly vulnerable, including the microbial communities supporting these biomes. Under extreme water deficit and high solar radiation, endolithic (inhabiting rock) habitats are considered environmental refuges for life (Friedmann, 1982; Cary et al., 2010; Pointing and Belnap, 2012; Wierzchos et al., 2012). In these microbial ecosystems, the rock substrate provides protection from incident UV, excessive solar radiation and freeze–thaw events while providing physical stability, and enhanced moisture availability (Walker and Pace, 2007; Chan et al., 2012). Endolithic microbial communities are photosynthetic-based with primary producers supporting a diversity of heterotrophic microorganisms (Walker and Pace, 2007; Wierzchos et al., 2012).

Habitats as different as the underside of quartz rocks (Cowan et al., 2011; Chan et al., 2013; de los Ríos et al., 2014), sandstones (Friedmann, 1982; de la Torre et al., 2003; Pointing et al., 2009), granite boulders (Wei et al., 2015), the inside of gypsum (Dong et al., 2007; Wierzchos et al., 2011; Wierzchos et al., 2015) and halite evaporites (Wierzchos et al., 2006; Robinson et al., 2015), carbonaceous (DiRuggiero et al., 2013) and volcanic rocks (Wierzchos et al., 2013; Cámara et al., 2014) among others, have shown that life has found innovative ways to adapt to the extreme conditions of hyper-arid deserts.

The Atacama Desert is one of the driest deserts on Earth and its hyper-arid core has been described as “the most barren region imaginable” (McKay et al., 2003). In the Yungay area of the hyperarid zone of the Atacama Desert, with decades between rainfall events and extremely low air relative humidity (mean yr^−1^ values <35%), ancient halite crusts of evaporitic origin have been shown to provide sufficient moisture to sustain cryptoendolithic (within pores) microbial communities (Wierzchos et al., 2006; de los Rios et al., 2010; Robinson et al., 2015). Another strategy for survival in this hyper arid desert is the colonization of cracks and fissures of rock substrates, also called chasmoendolithic colonization (DiRuggiero et al., 2013). Our work in the hyper-arid core of the Atacama Desert revealed that such communities inhabit gypsum covered rhyolite and calcite rocks, where the complex network of cracks and fissures of the rocks promote water retention (DiRuggiero et al., 2013). Calcite is highly translucent, with a network of large connected fissures providing abundant space for microbial colonization within a few cm of the rock surface (DiRuggiero et al., 2013). We found that the diversity of the calcite community was significantly increased when the water budget of the ecosystem was potentially enhanced by dew formation on the rock in contrast to the rhyolite community, where the unique source of water were scarce rainfalls (DiRuggiero et al., 2013). More recently, we discovered communities colonizing ignimbrite, a pyroclastic rock composed of plagioclase and biotite embedded in a matrix of weakly-welded glass shards (Wierzchos et al., 2013). In the ignimbrite the colonization is cryptoendolithic, with microorganisms colonizing small pores within the rock and only within a few mm of the rock surface (Wierzchos et al., 2013; Cámara et al., 2014).

From previous work on microbial communities in hyper arid deserts, it is clear that the climate regime play a major role in determining the habitability of a given substrate. However, the susceptibility of rocks to colonization also depends on the architecture of the lithic substrate (Wierzchos et al., 2015), including factors such as translucence, which allows transmission of photosynthetically active radiation (PAR), thermal conductivity, the presence of a network of pores and/or fissures connected to the rock surface, which is linked to the capacity at retaining water, and chemical composition. These factors are likely to impact greatly microbial colonization and diversity (Herrera et al., 2009; Cockell et al., 2011; Walker and Pace, 2007; Wierzchos et al., 2012).

Nevertheless, very little information is available about what rock properties are driving endolithic communities functioning and diversity. Here we used a metagenomic analysis of two types of endolithic colonization in calcite (chasmoendolithic) and ignimbrite (cryptoendolithic) rocks collected in the extreme environment of the hyper-arid core of the Atacama Desert. The taxonomic and functional composition of the communities were characterized and interpreted within the context of the structural properties of each rock

## Material and Methods

### Sampling and site characterization

Calcite (composed mainly of calcium carbonate) rocks were collected in 2013 near Valle de la Luna (VL and CB) (DiRuggiero, 2013) and ignimbrite (pyroclastic volcanic glass) rocks were collected in the Lomas de Tilocalar area (IG and MIG) (Wierzchos et al., 2013) of the Atacama Desert in northern Chile. Additional ignimbrite rocks were also collected from a nearby area 25 km to the North-East of Lomas to Tilocalar (PING). Microclimate data for the Lomas de Tilocalar area was collected *in situ* from April 2011 to 2013 using an Onset HOBO^®^ Weather Station Data Logger (H21–001) as previously described (Wierzchos et al., 2013). Solar flux was measured using a PAR sensor for wavelengths of 400–700 nm (Wierzchos et al., 2013). Microclimate data for Valle de la Luna was extracted from historical records for the village of San Pedro de Atacama located 5 km east of our sampling site, as reported by DiRuggiero et al. (DiRuggiero et al., 2013).

### Microscopy analyses

Colonized rock samples were processed for scanning electron microscopy in backscattered electron mode (SEM-BSE) observation and/or energy dispersive X-ray spectroscopy (EDS) microanalysis according to methods by Wierzchos et al. (Wierzchos and Ascaso, 1994; Wierzchos et al., 2011). SEM-BSE was used in combination with EDS to characterize the lithic substrates. Rock samples were observed using a scanning electron microscope (DSM960 Zeiss; Carl Zeiss) equipped with a solid-state, four diodes BSE detector plus an auxiliary X-ray EDS microanalytical system (Link ISIS Oxford, UK).

Fluorescence microscopy (FM) in structural illumination microscopy mode (SIM) using DAPI nucleic acids stain was performed on cell aggregates gently isolated from the chasmoendolithic habitat (Wierzchos et al., 2011). The samples were examined using a fluorescence microscope (AxioImager M2, Carl Zeiss, Germany) in SIM mode with a ApoTome (commercial SIM by Zeiss) system for 3 dimensional (3D) visualization of cell aggregates (Wierzchos et al., 2011).

### DNA extraction and sequencing

For 16S rRNA gene sequencing, total genomic DNA was extracted from rock powder using the PowerSoil DNA Isolation kit (MoBio Laboratories Inc., Solana Beach, CA) from four calcite and four ignimbrite rocks. DNA was amplified using the barcoded universal primers 338F and 806R for the V3–V4 hypervariable region. Amplicons from at least 3 amplification reactions were pooled together, purified, and sequenced using the Illumina MiSeq platform by the Genomics Resource Center (GRC) at the Institute for Genome Sciences (IGS), University of Maryland School of Medicine. For metagenomes, the DNA was extracted from 4 rocks for each substrate from above (VLC and IG), pooled, and sequencing libraries were prepared using the Nextera XT DNA sample preparation kit (Illumina, San Diego, CA), with an average insert size of 400 bp, and sequencing was performed on the Illumina HiSeq2500 platform by the GRC at IGS.

### 16S rRNA gene sequences analysis

Illumina paired-end sequences were processed using the QIIME package (v1.6.0) (Caporaso et al., 2010) as previously described (DiRuggiero et al., 2013). Alpha and beta diversity metrics were calculated based on OTUs at the 0.03% cutoff (OTUs0.03) in QIIME. Statistical testing using Nonparametric Mann-Whitney tests were performed to indicate confidence in similarities/differences observed.

### Metagenome analysis and assembly

Illumina sequencing of the metagenomic libraries produced 126,820,576 and 100,086,740 paired-end reads for the calcite and ignimbrite samples, respectively. Ribosomal RNA sequence reads were removed using Bowtie (v1) (Langmead et al., 2009) and by mapping to SILVA reference database (Pruesse et al., 2007). Both paired-end reads were filtered out if one read mapped to the rRNA database. Raw reads with low-quality bases as determined by based calling (phred quality score of 20 that corresponds to an error probability of 1%) were trimmed from the end of the sequence, and the sequence longer than 75% of the original read length were retained. The reads were assigned taxonomy content using PhyloSift (Darling et al., 2014) and functional content using HUMAnN (Abubucker et al., 2012) and the Metagenomics analysis server MG-RAST (Meyer et al., 2008). An assembly was produced for the metagenome using the IDBA-UD assembler for metagenomic sequencing data (Peng et al., 2012). We use a *k* range of 20–100 with a pre-correction step before assembly. Assembled contigs were grouped into potential draft genomes using MaxBin (Wu et al., 2014), an expectation-maximization (EM) algorithm using tetranucleotide frequencies, abundance levels, and single-copy gene analysis. Taxonomy was assigned to each bin using both PhyloSift and Kraken (Darling et al., 2014; Wood and Salzberg, 2014).

### Phylogenetic Analysis

The phylogenetic position of cyanobacteria metagenomic bins was determined using marker genes extracted by Phylosift and that were conserved across metagenomic bins with at least 50% estimated genome completeness from MaxBin (Wu et al., 2014). The marker genes were aligned individually to reference genes from the UniProt database using MUSCLE (Edgar, 2004) and the individual alignments were concatenated. Low-quality regions of the alignment were removed using Gblocks (Castresana, 2000) and a maximum-likelihood phylogenetic tree was created using FastTree (Price et al., 2010).

### Population genome annotation

Cyanobacterial genomic bins Ca9 and Ig12 were selected for annotation as draft genomes based on both estimated genome completeness and n50 values of the binned contigs. RAST server (Aziz et al., 2008) was used to annotate and compare the draft genomes to known references. Putative phage contigs incorrectly binned were identified using the annotation and removed from the draft genomes. The antibiotics and Secondary Metabolite Analysis Shell antiSMASH 2 (Blin et al., 2013) was used for the prediction of non-ribosomal peptides (NRPS) structures and synthetic gene clusters. The NORINE database (http://bioinfo.lifl.fr/norine/) was used to determine chemical structures for putative products of NRPS and synthetic gene clusters. The draft genomes were compared using protein products predicted with Prodigal (Hyatt et al., 2010) and BLASTP (Gish and States, 1993) with a cutoff of 80% amino acid identity between matches. Results from the draft genomes comparisons were visualized using Circos (Krzywinski et al., 2009).

### Sequence data and availability

All sequences were deposited at the National Center for Biotechnology Information Sequence Read Archive under Bioproject ID PRJNA285514 and accession numbers SAMN03754650 (ignimbrite) and SAMN03754649 (Calcite). The MG-RAST report for the data is available under ID 4600829.3 and 4600830.3 (http://metagenomics.anl.gov/metagenomics.cgi?page=MetagenomeOverview&metagenome=4600829.3 and http://metagenomics.anl.gov/metagenomics.cgi?page=MetagenomeOverview&metagenome=4600830.3). Completed assemblies, annotation, and phylogenetic trees are available at http://figshare.com/articles/Phylogenetic_and_Functional_Substrate_Specificity_for_Endolithic_Microbial_Communities_from_the_Atacama_Desert/1606253.

## Results

We characterized at the molecular level the endolithic microbial communities from two rock types from the Atacama Desert, calcite from Valle de La Luna and ignimbrite from Lomas de Tilocalar. Both areas are extremely dry with less than ~ 25 mm rainfall each year and average air RH between 18.3 and 40.5%, (Table 1). In both locales, temperatures reached values above 32°C and maximum PAR values were about 2500 μmol s^−1^m^−2^.

**Table 1:** Microclimate data for the VL (calcite) and LdT (ignimbrite) sampling sites

| Sampling sites | PAR $\mu\text{mol m}^{-2} \text{s}^{-1}$ | | Air temperature ($^{\circ}\text{C}$) | | | Air relative humidity (%) | | | Rainfall (mm) |
| --- | --- | --- | --- | --- | --- | --- | --- | --- | --- |
|  | ave | max. | min. | ave | max. | min. | ave | max. | ave |
| Valle de la Luna (VL)<sup>a</sup> | - | 2600 | -1.8 | 13.1 | 32.2 | 16.7 | 40.5 | 80 | 27.8 |
| Lomas to Tilocalar (LdT)<sup>b</sup> | 442 | 2553 | -4.6 | 15 | 44.8 | 1.2 | 18.3 | 100 | 25.6 |

Maximum (max.), minimum (min.) and yearly average (ave) values for photosynthetic active radiation (PAR), temperature, and relative humidity (RH); <sup>a</sup>data from DiRuggiero et al., 2013; <sup>b</sup>data collected by HOBO weather station from April 2011-2013; “-” no data.

The calcite samples were composed of laminated calcite layers with a thickness of several centimeters. The calcite mineralogical composition and petrographic study were previously reported by DiRuggiero et al. (2013). The calcite rock surface was covered by a hardened semitransparent layer (up to 5 mm thick) with microrills forming small and short sinuous channels on the rock surface (Fig. 1a). The most characteristic feature of the calcium carbonate rock was the presence of irregular narrow fissures and cracks that extended perpendicular and/or parallel to the rock surface (Fig. 1a and 1b). A large number of these fissures and cracks, up to dozens of millimeters in length, were colonized by chasmoendolithic microorganisms detected upon fracturing the rock (Fig. 1a). Additional fractures, parallel to the rock surface, also revealed similar chasmoendolithic colonization (black arrow on Fig. 1a). Using SEM-BSE, we observed an almost continuous colonization of these fissures by chasmoendoliths (Figs 1b-e) and the connection of the fissures to the rock surface (Figs 1). We observed mostly well-developed cyanobacteria cells surrounded by one or two sheaths of capsular-like extracellular polysaccharidic substances (EPSs) and heterotrophic bacteria present within some of the cyanobacteria aggregates (arrow in Fig. 1d). Closer to the rock surface, cyanobacteria cells appeared deteriorated with collapsed cells and remains of microbial colonization (arrows in Fig. 1e). The 3-Dimensional reconstruction of the cyanobacteria aggregates shown characteristic microbial aggregates inside sack-like structures formed by EPSs (green signal in Fig. 1f).

**Fig. 1.**
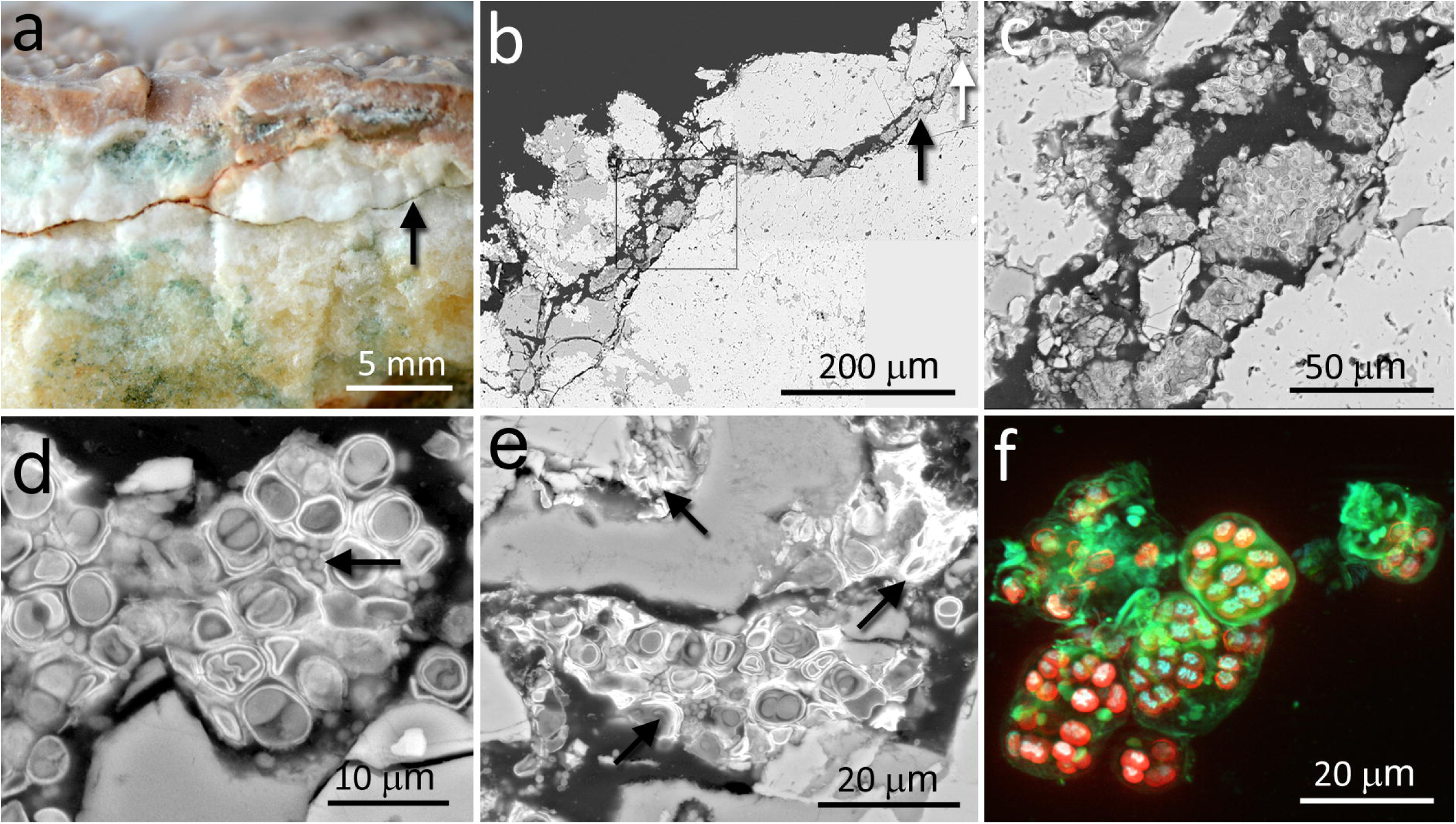
Chasmoendolithic microbial communities within the fissures and cracks of calcite rock. (a) Freshly fractured calcite show fissure wall with green spots – chasmoendoliths; black arrow point to parallel to the rock surface fissure also colonized; white arrow, hardened layer with microrills (asterisk). (b) SEM-BSE low magnification image of transversal section showing the connected to the rock surface fissure full-fill by chasmoendolithic colonization; the square, black and white arrows point to detailed (c), (d) and (e) images respectively. (c) SEM-BSE image shows chasmoendolith’s aggregates colonizing fissure walls. (d) SEM-BSE image shows well developed cyanobacteria surrounded by EPSs and heterotrophic bacteria (arrow). (e) SEM-BSE image shows collapsed cyanobacteria cells and remains of microbial colonization (black arrows). (f) SIM 3D image of cyanobacteria aggregates; red signal - cyanobacteria photosynthetic pigments autofluorescence, blue signal - cyanobacteria nucleoids and green signal sheaths of EPSs surrounding cyanobacteria aggregates and singular cells.

In contrast to the chasmoendolithic habitat found within the calcite fissures, the cryptoendolithic colonization within the volcanic ignimbrite presented a very different picture. The ignimbrite has a structure of vitrified foam identical to pumice rock with a large number of connected and non-connected bottle-shaped pores. The mineralogical, petrographic, and structural characteristics of the ignimbrite were previously reported by Wierzchos et al. (Wierzchos et al., 2013). The colonization zone of ignimbrite rocks was in the form of a narrow (1–3 mm thick) intensively green layer, parallel to the ignimbrite surface (Fig. 2a), and within 2–3 mm of the rock surface. This brown-colored varnish rock covering the surface was composed of allochthonous clay minerals and filling some of the pores at the surface (white arrow in Fig. 2b). Just below the ignimbrite surface cover, a cryptoendolithic colonization zone was observed (artificially cyan-colored spots in Fig. 2b, black arrow) with many small pores. The microbial assemblage within the bottle-shaped pores was composed of cyanobacteria and heterotrophic bacterial cells, seemingly not associated with EPSs (Fig. 2c-d). Some of the pores were not colonized whereas others were almost totally filled by cryptoendoliths (Fig. 2d). Cyanobacteria formed aggregates and, in places, cells were associated with heterotrophic bacteria (arrow in Fig. 2e). In some pores, we also observed both damaged in healthy cyanobacteria together with heterotrophic bacteria, probably also damaged (Fig. 2f).

**Fig. 2.**
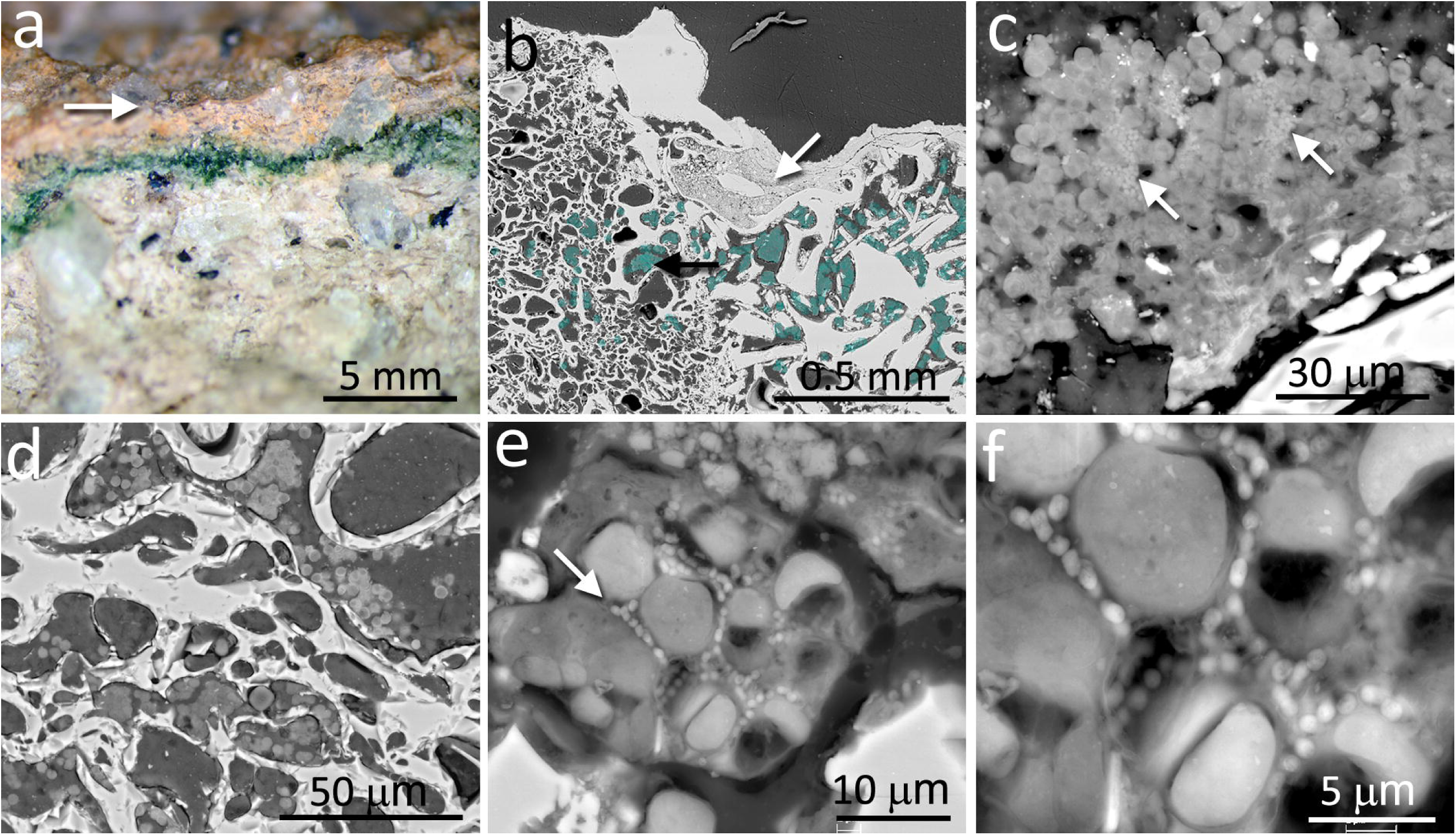
Cryptoendolithic microbial communities within the pores of ignimbrite. (a) Freshly fractured ignimbrite show narrow green layer of cryptoendolithic microorganisms beneath the rock surface (arrow). (b) SEM-BSE image showing micromorphological features of rhyolitic ignimbrite composed of glass shards welded together forming bottle-shaped vesicles (left image side) and irregular-shape pores (right image side); white arrow point to the surface pore filled by varnish rock mineral components; aggregates of cryptoendolithic microbial colonization were marked by cyan-colour and black arrow points to aggregate in detail shown in fig. 2c. (c) SEM-BSE image of cyanobacteria composed aggregate containing heterotrophic bacteria (white arrows). (d) SEM-BSE image of empty and colonized bottle-shaped pores. (e) SEM-BSE image of detail view of cyanobacteria aggregates containing heterotrophic bacteria (white arrow). (f) Magnified central part of fig. 2e showing heterotrophic bacteria in close position to the declined cyanobacteria and their remains.

### Taxonomic diversity of the rock communities

We obtained a total of 315,334 16S rRNA gene paired-end sequences, with an average read length of 408 bp, for four individual rocks from each of the two endolithic substrates, calcite and ignimbrite. Taxonomic assignments of the sequence reads revealed that the dominant phyla, in both samples, were *Cyanobacteria, Actinobacteria*, and *Chloroflexi*, and that their relative distribution was significantly different between the two substrates (Fig. S1). Diversity metrics at a maximum sequencing depth of 2400 sequences per sample support the finding that the calcite substrate harbors a more diverse community in term of richness and phylogenetic diversity (Table S1; Fig. S2). Nonparametric Mann-Whitney tests reported that this difference is significant (p<0.05) across all diversity metrics. Although the statistical power of this test is weak due to the small sample size of this comparison (n=8), the ranges for each substrate of all diversity metrics are non-overlapping and distinct (Table S1). Photosynthetic cyanobacteria dominated both communities and greater proportional abundances were found in all the ignimbrite samples than in the calcite samples; this was also supported by a nonparametric Mann-Whitney test (p=0.03). Fourteen and 12 major OTUs of cyanobacteria (>1% abundance) were identified in the calcite and ignimbrite communities, respectively, and none were shared across the two communities.

We generated over 22 Gbp of high quality, paired-end, metagenomic shotgun sequences for the calcite and ignimbrite substrates (Table 2). Taxonomic assignment of the metagenomic reads was performed with PhyloSift using all extracted marker genes (Fig. 3). When compared to the 16S rRNA gene taxonomic distribution, averaged for each rock substrate, we found a general consensus between the two datasets (Fig. 3), which supported a higher taxonomic diversity for the calcite community. In both substrates the metagenome analysis showed higher proportional abundance of rare taxa, probably as the result of variation in genome sizes across all taxa. Reconstruction of 16S rRNA gene sequences from the metagenomic dataset with EMIRGE (Miller et al., 2011) provided full-length genes for all the major phyla observed in the 16S rRNA gene dataset (Fig. S3) and indicated again a higher taxonomic diversity in the calcite community.

**Table 2:** Number of sequence reads, Gbp of sequences, and contigs statistics for the calcite and ignimbrite metagenomes

| Sample | Number of paired-end reads<sup>(1)</sup> | Number of Gbp sequenced | Number of Gbp assembled | Number of contigs | Contig mean length (bp) | Contig maximum length (bp) |
| --- | --- | --- | --- | --- | --- | --- |
| Ignimbrite (MTQ) | 100,086,740 | 10.1 | 0.22 | 166,509 | 1,351 | 101,619 |
| Calcite (VL) | 126,820,576 | 12.8 | 0.35 | 344,986 | 1,016 | 119,637 |

<sup>(1)</sup>after quality control

**Fig. 3.**
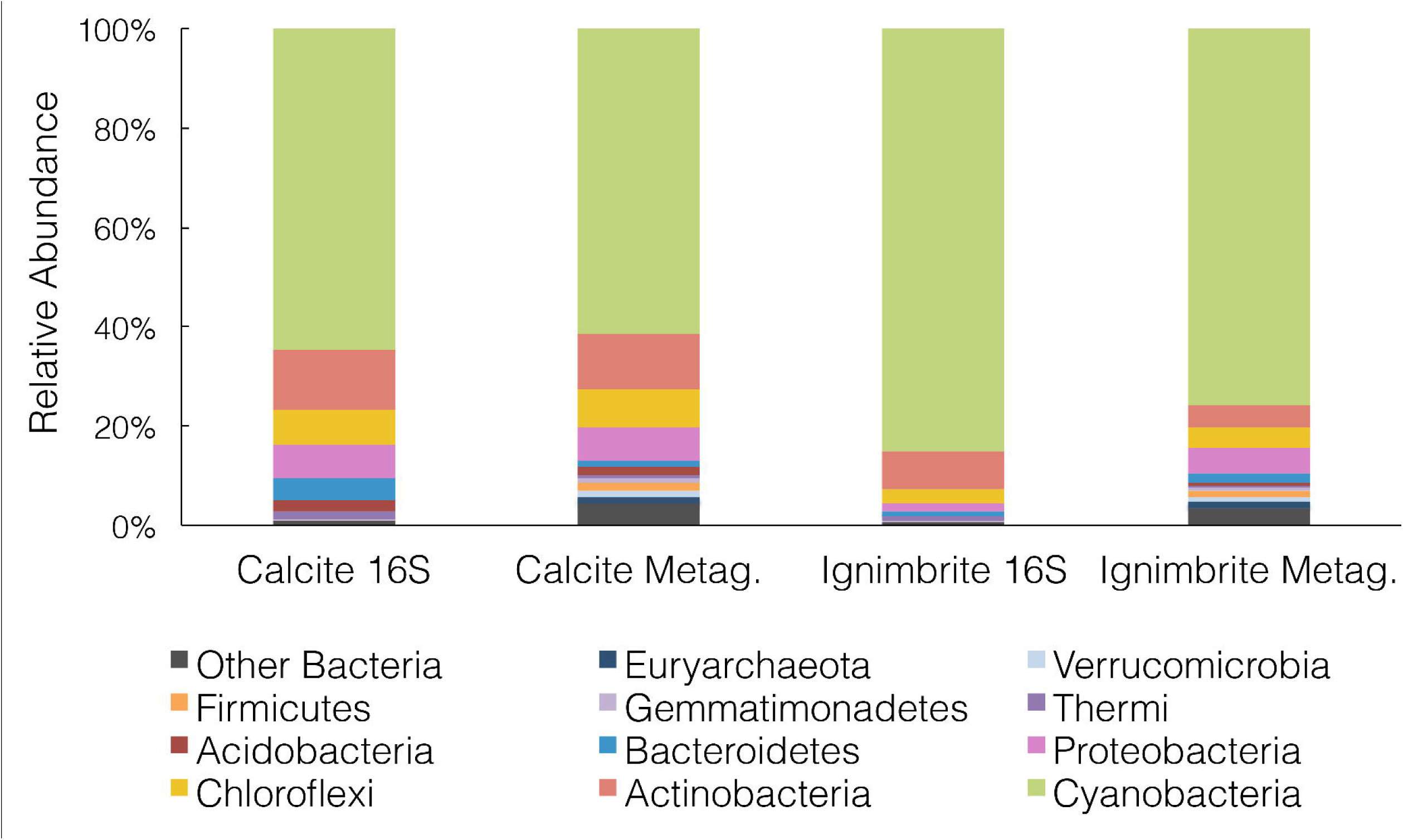
Taxonomic composition at the phylum level for the calcite and ignimbrite communities using either 16S rRNA gene sequences (16S) or PhyloSift marker genes from the metagenome data (Metag) (10% relative abundance cut-off).

### Functional diversity of the rock communities

The functional composition of the calcite and ignimbrite metagenomes was analyzed with HUMAnN and MG-RAST using total sequence reads. The HUMAnN analysis showed that at the KEGG Super-pathway level, the two communities maintained a strong degree of similarity in functional composition (Fig. S4). Functional assignment of sequence reads to the SEED or COG database by MG-RAST (Fig. 4) also showed high similarity between the two communities. When using the SEED database to classify sequence reads into metabolic functions, categories with the most reads were for metabolisms related to “carbohydrates”, “proteins”, “amino acids and derivatives”, and “cofactors, vitamins, prosthetic groups and pigments” (Fig. 4).

**Fig. 4.**
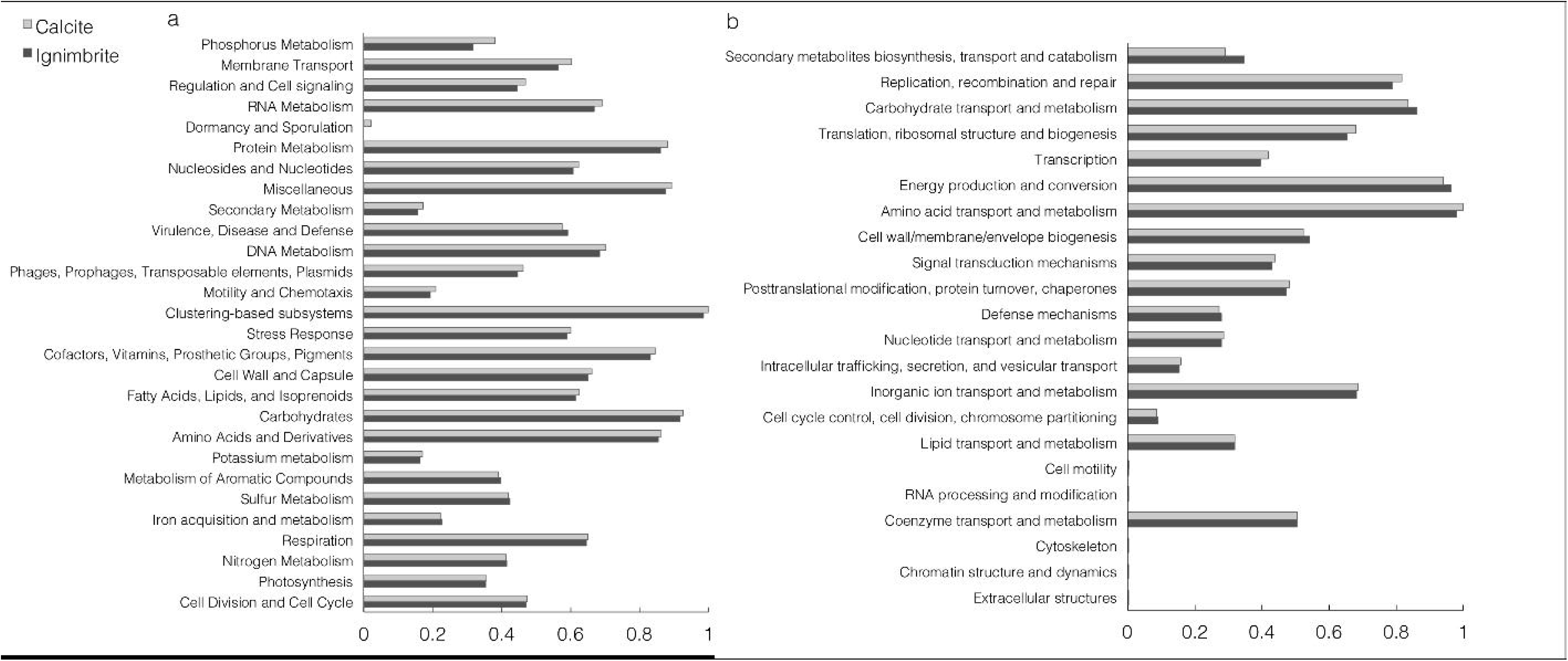
Normalized abundance of functional categories for the calcite and ignimbrite community metagenome sequence reads for (a) SEED subsystems and (b) COG generated with MG-RAST.

**Fig. 5.**
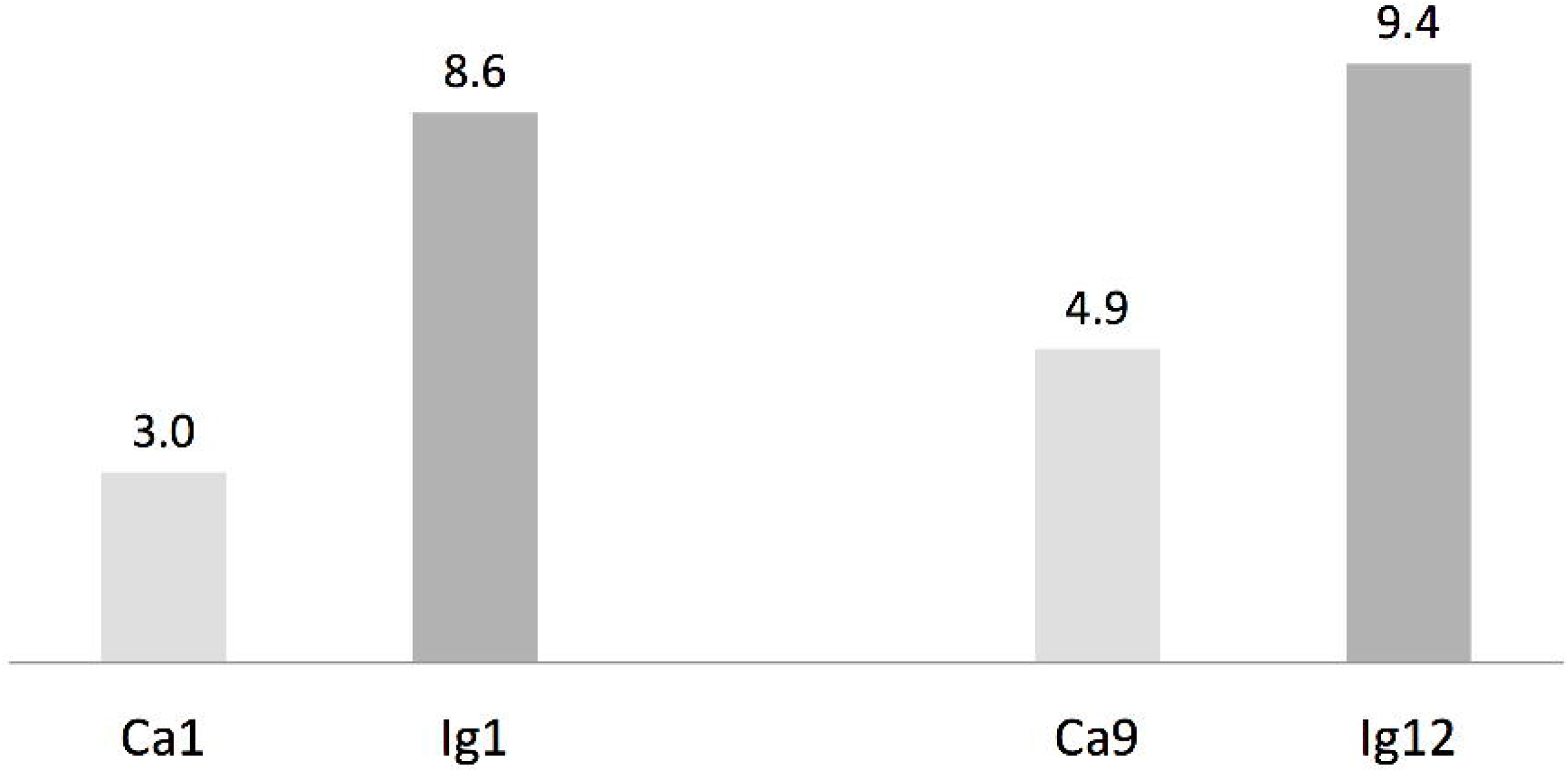
Number of non-ribosomal peptides and polyketides genes per Mbp in cyanobacteria draft genomes from the calcite (Ca1 and Ca9) and ignimbrite (Ig1 and Ig12) communities using AntiSmash predictions.

The metabolic potential for both the calcite and ignimbrite communities was further analyzed using the MG-RAST server. Genes encoding for pathways for photosynthesis and auxotrophic CO_2_ fixation were found in both the calcite and ignimbrite metagenomes and included biosynthetic genes for photosystems I and II (PSI and PSII), phycobilisome (light harvesting antennae of PSII in cyanobacteria), the Calvin-Benson cycle, carboxyzomes, and photorespiration. Genes for nitrogen fixation were not detected, despite an intensive search using BLASTP and HMMER (Finn et al., 2011) on all predicted proteins from the sequence reads and from the assembled metagenomes. Using the SEED functional classification, we found that the major pathways for nitrogen acquisition in both communities were ammonia assimilation (direct uptake of ammonia and assimilation into glutamine and glutamate) and nitrate and nitrite ammonification (direct uptake of nitrate or nitrite and subsequent reduction to ammonia); minor pathways included allantoin utilization and cyanate hydrolysis (Fig. S5).

Two major pathways for phosphorus acquisition were identified in both communities from the abundance of their protein encoding genes. One extensive suite of genes constituted a putative Pho regulon and another set of genes encoded for proteins involved in the transport and metabolism of phosphonates. With respect to iron transport, the calcite and ignimbrite metagenomes contained mostly siderophore/heme iron transporters. These ABC transporters are known to mediate the translocation of iron-siderophore complexes across the cytoplasmic membrane (Köster, 2001). We also found ABC transporters of the ferric iron type and, to a lesser extent, of the ferrous iron type and known to be induced by low pH (Köster, 2001).

The functional analysis of the metagenomes from both communities also uncovered a diversity of stress response pathways, with most notably genes involved in carbon starvation, cold shock, oxidative stress, osmotic stress/desiccation tolerance, and secondary metabolites. In both the calcite and ignimbrite metagenomes, we found genes encoding for polyol transport (ABC transport systems), in particular glycerol, choline, betaine, and glycine betaine uptake systems (ABC transport systems), betaine and ectoine biosynthesis, and synthesis of glucans (Table S3). A gene encoding an aquaporin Z water channel, mediating rapid flux of water across the cellular membrane in response to abrupt changes in osmotic pressure (Calamita, 2000), was also found in high abundance. Also notable, the large number of genes dedicated to trehalose biosynthesis and uptake (Table S3). Secondary metabolites included a wide variety of compounds including non-ribosomally made peptides (NRPS), polyketides (PKS), and alkaloids. The majority of genes encoding for NRPS and PKS represented a wide range of siderophores, including pyoverdine, pyochelin, and yersiniabactin (Table S2). Biosynthetic pathways for other secondary metabolites included those for alkaloids, phenylpropanoids, phenazines, and phytoalexins.

The two SEED subsystems of greatest difference between the two communities were phosphorous metabolism and membrane transport, with both functional categories showing higher gene abundance in the calcite community (Fig. 4a). Functional diversity was also higher in the calcite community for genes involved in iron acquisition and transport, Mn transport, and the synthesis of compatible solutes such as betaine, ectoine, and glucans (Table S3). One notable exception was a ferrichrome iron receptor, which was about 3 times more abundant in the ignimbrite than in the calcite community (Table S3); the encoding genes were all annotated to cyanobacteria taxa. Overall, the pathways involved in osmoregulation were more diverse in the calcite than in the ignimbrite community (Table S3). In addition, the functional categories most enriched in the calcite community were not from the most abundant taxa; for example, genes involved in the synthesis of osmoregulated periplasmic glucans, and most genes for the ectoine biosynthesis pathways, were from *Proteobacteria, Actinomycetales*, and *Firmicutes* rather than from *Cyanobacteria* or *Rubrobacter*. In contrast, when looking at gene abundance rather than diversity, we found considerably more NRPS and PKS related genes, especially siderophores, in the ignimbrite metagenome (Table S2). The COG category for secondary metabolite biosynthesis/transport also shows similar higher levels of these genes in the ignimbrite metagenome (Fig. 4b).

### Sequence assembly

The 12.8 Gbp of metagenome sequence data for the calcite community was assembled into a total of 350 Mbp of contigs (344,986) with an n50 value of 1,530 bp and a maximum contig length of 101,619 bp. The 10.1 Gbp metagenome sequence data for the ignimbrite community were assembled into 220 Mbp of contigs (166,509) with an n50 value of 2,631 bp and a maximum length of 119,637 bp (Table 2). Binning of contigs into population genomes using MaxBin generated 42 and 34 putative population genome bins, for the calcite and ignimbrite communities, respectively. All bins were individually assigned taxonomy using both Kraken and PhyloSift, and bins mostly composed of single taxa were further analyzed (Table 3). We found that the bins’ G+C content closely matched that of the reference species, with G+G below 50% for *Cyanobacteria* and above 65% for *Rubrobacter* species. Interestingly, the relative abundance ratio of a bin did not necessarily match its estimated completeness. Rather, population genomes with the highest coverage produced lower quality assemblies, i.e. more contigs of smaller sizes (compare for example Ca01 with Ca09, and Ig01 with Ig12). This likely resulted from a higher representation of micro-heterogeneity at high coverage, for a given population genome, impairing the assembler functionality.

**Table 3:** Most abundant species in the calcite and ignimbrite communities obtained by binning the metagenome sequence data

| Calcite |  |  |  |  |  |  | Ignimbrite |  |  |  |  |  |  |
| --- | --- | --- | --- | --- | --- | --- | --- | --- | --- | --- | --- | --- | --- |
| Bin number | Taxa | %<sup>(a)</sup> | Estimated completeness % | Number of contigs | Size Mbp | G+C % | Bin number | Taxa | %<sup>(a)</sup> | Estimated completeness % | Number of contigs | Size Mbp | G+C % |
| Ca01 | *Gloeocapsa sp.* | 20.7 | 95.3 | 2329 | 7.7 | 48 | lg01 | *Chroococcidiopsis sp.* | 32.8 | 73.8 | 2031 | 5.1 | 45 |
| Ca02 | *Gloeocapsa sp.* | 9.2 | 46.7 | 2388 | 5.6 | 46 | lg02 | *Rubrobacter xylanophilus* | 15.1 | 38.3 | 1215 | 2.2 | 68 |
| Ca03 | *Rubrobacter xylanophilus* | 4.2 | 43 | 413 | 1.2 | 65 | lg04 | *Rubrobacter xylanophilus* | 5.9 | 31.8 | 1900 | 6 | 67 |
| Ca04 | *Rubrobacter xylanophilus* | 3.9 | 35.5 | 330 | 1.5 | 69 | lg05 | *Frankineae sp.* | 5.1 | 74.8 | 460 | 4.5 | 75 |
| Ca05 | *Acaryochloris sp.* | 3.6 | 68.2 | 1027 | 5.5 | 48 | lg06 | *Conexibacter woesei* | 2.7 | 87.9 | 357 | 3.9 | 75 |
| Ca06 | *Conexibacter woesei* | 3.4 | 45.8 | 307 | 1.9 | 74 | lg07 | *Frankineae sp.* | 2.3 | 93.5 | 457 | 3.9 | 72 |
| Ca07 | *Thermomicrobia sp.* | 3.2 | 83.2 | 1161 | 3.9 | 72 | lg09 | *Thermobaculum sp.* | 2.0 | 86 | 661 | 3.5 | 69 |
| Ca09 | *Gloeocapsa sp.* | 2.8 | 94.4 | 800 | 6.5 | 45 | lg10 | *Thermomicrobia sp.* | 2.0 | 89.7 | 548 | 5.1 | 63 |
| Ca17 | *Conexibacter woesei* | 1.9 | 86 | 483 | 3.3 | 72 | lg11 | *Conexibacter woesei* | 1.9 | 59.8 | 575 | 2.9 | 76 |
| Ca19 | *Microlunatus sp.* | 1.6 | 91.6 | 538 | 4.3 | 71 | lg12 | *Chroococcidiopsis sp.* | 1.7 | 93.5 | 817 | 6.6 | 46 |

<sup>a</sup>Relative abundance ratios; defined as actual coverage levels divided by summed coverage levels of all genomes.

The most abundant population genomes found in the calcite and ignimbrite metagenomes were related to *Chroococcidiopsis* and *Gloeocapsa*, both members of the *Chroococcales*. Cyanobacteria of the *Acaryochloris* genus were found only in the calcite community. The *Actinobacteria* were dominated by *Rubrobacter xylanophilus*, together with *Conexibacter woesei* and a number of other *Actinomycetales* species, the later differing between the two communities. *Chloroflexi* populations composed of *Thermomicrobia*, *Chloroflexus*, and *Thermobaculum* species were also found in the ignimbrite and calcite communities. The dominant *Proteobacteria* was *Sphingomonas*, and in the ignimbrite community, the low G+C% *Bacteroidetes* species *Segetibacter* was represented (Table 3). The phylogenetic analysis of the metagenomic bins indicated that there was significantly greater diversity within the calcite *Cyanobacteria* community, with bins distributed in two distinct clades, whereas only one *Cyanobacteria* clade was found within the ignimbrite dataset (Fig. S6). The ignimbrite *Cyanobacteria* population genomes were distinct at the phylogenetic level to that of the calcite, but one of the calcite metagenomic bins (Ca02) was consistently more closely related to the ignimbrite than the others calcite bins, further supporting the extent of the taxonomic diversity of the calcite community (Fig. S6).

A candidate draft genome for the dominant *Cyanobacteria* population genome in each community was selected based on estimated completeness, calculated from single gene copy analysis (MaxBin). The estimated completeness was 94.4% for *Gloeocapsa sp*. Ca09, from the calcite metagenome, and 93.5% for *Chroococcidiopsis sp*. Ig12, from the ignimbrite metagenome (Table 3). The genomic bins were annotated using the RAST server, and the two genomes were found to share similar hits for 64% of their total gene products. Three contigs from Ca09 and five contigs from Ig12 were found to be entirely composed of phage-related and hypothetical proteins, making them likely to be cyanophage genomic fragments binned together with the host genome. This co-binning was probably due to the tendency of bacteriophages to share tetranucleotide frequencies, genes, and codon usage patterns with their host (Soueidan et al., 2014). These contigs were removed from the draft genomes for the analysis of annotation features and gene products. On average the two annotated proteomes had 69.4% amino acid identity, and only 27% and 24% of the gene products could be assigned a SEED category for Ca09 and Ig12, respectively. The draft genomes had multiple genes involved in choline and betaine biosynthesis subsystem and Aquaporin Z, both related to osmotic stress. A Mycosporine synthesis cluster, MysA, MysB, and MysC, was found in the Ca09 but not in Ig12 draft genome; a BLAST search against the entire metagenomes found these genes to be absent from the ignimbrite metagenome. The Ca09 draft genome had six copies of the ferrichrome-iron receptor and the Ig12 draft genome had 12, while the closest reference genome in RAST, *Nostoc punctiforme PCC 73102*, only had one copy of this gene. Another iron related gene, the Iron(III) dicitrate transport system FecB gene, had multiple duplications in both Ca09 and Ig12, but is absent in the *Nostoc punctiforme* RAST annotation.

We used antiSMASH to identify biosynthesis loci for secondary metabolism in the four major cyanobacteria population genomes from the calcite and ignimbrite metagenomes (Table 3). Similarly to the whole metagenome, we found an increased abundance of NRPS and PKS secondary metabolite genes and gene clusters in the Ig12 ignimbrite genome. Searches in the NORINE database (http://bioinfo.lifl.fr/norine/) revealed that the closest chemical structures for the putative products of these gene clusters were often siderophores (Fig. 6). A figure linking genes products with greater than 80% amino acid identity between the complement of contigs for the Ca9 and Ig12 draft genomes illustrates the close relatedness between these strains and also the greater abundance of NRPS/PKS and Ferrichrome-iron receptor proteins in the draft genome from the ignimbrite community (Fig. 7).

**Fig. 6.**
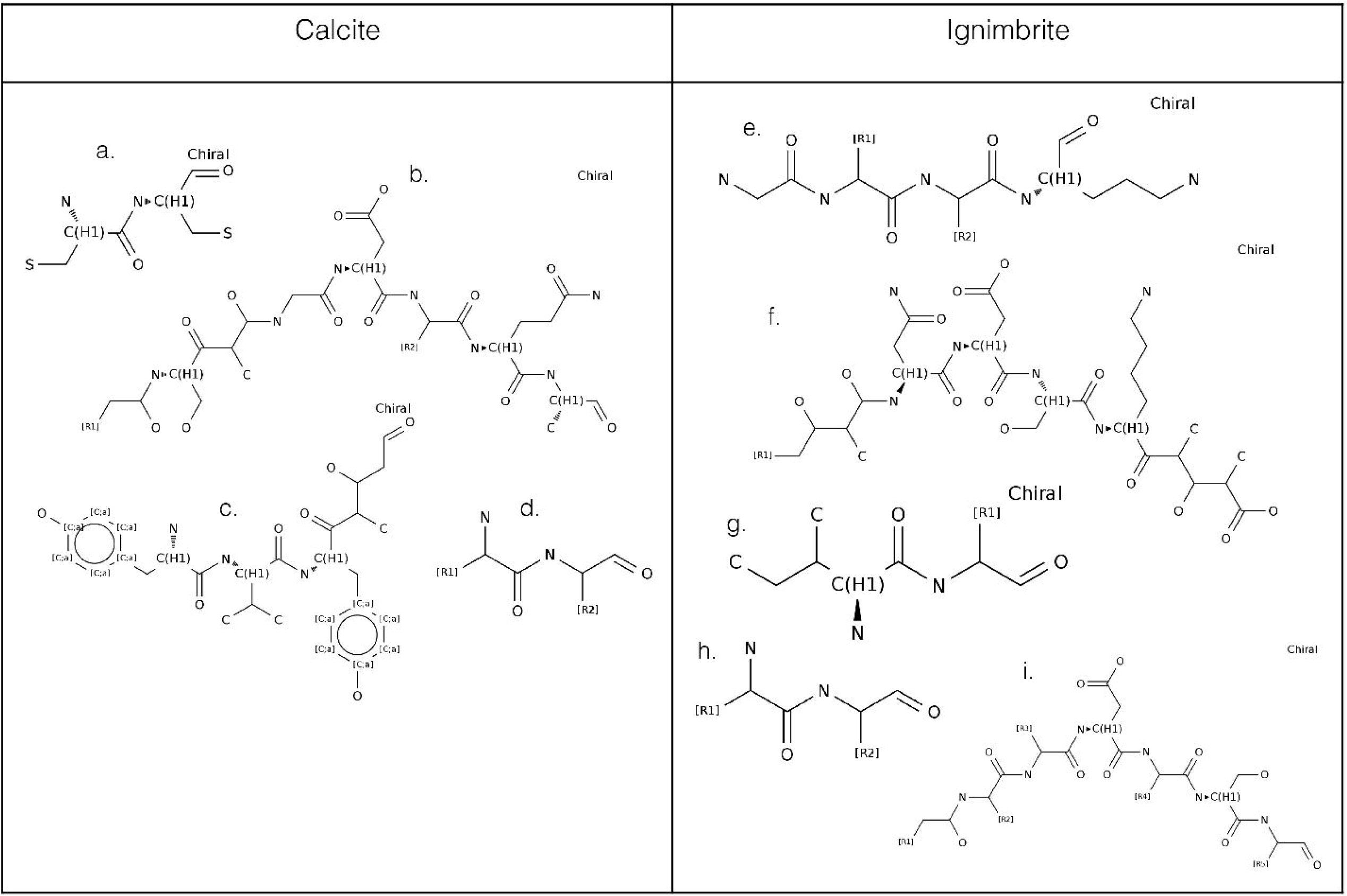
NORINE predictions for putative products of non-ribosomal peptides and polyketides gene clusters found in the draft genomes of cyanobacteria from the calcite (Ca9) and ignimbrite (Ig12) communities.

**Fig. 7.**
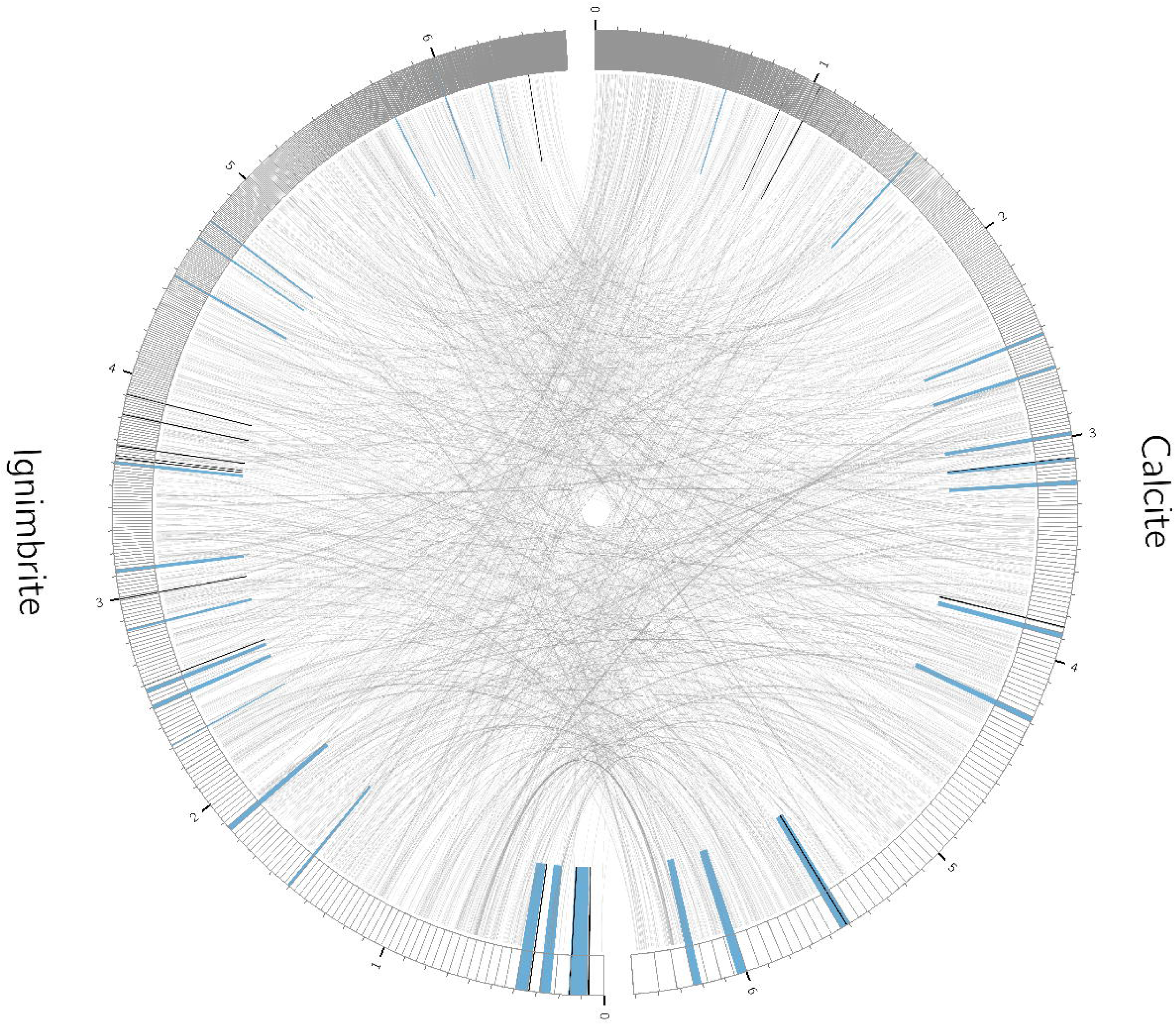
Relationships between contigs from the calcite Ca09 and ignimbrite Ig12 draft genomes. Links are between gene products with greater than 80% amino acid identity. Highlighted in blue are secondary metabolite clusters identified by antiSMASH, and marked in black are Ferrichrome-iron receptor proteins. Figure produced with Circos (Krzywinski 2009).

## Discussion

By linking taxonomy to function, our study of endolithic systems in the Atacama Desert provides an understanding of microbial adaptation strategies to extreme desiccation and solar irradiance. It also provides an integrated view of the biotic and abiotic factors shaping the functioning of these highly specific ecosystems. Functional metagenomic analysis was carried out on two rock substrates, calcite and ignimbrite, and the context for the interpretation of the molecular data was provided by the examination of water availability, the mineralogical composition, and the type of colonization (chasmo-versus cryptoendolithic) for the two types of rock.

The Valle de la Luna (calcite rocks) and Lomas de Tilocalar (ignimbrite rocks) sampling sites are located in the hyper-arid zone of the Atacama Desert and, as such, both locations experience rare precipitations (DiRuggiero et al., 2013; Wierzchos et al., 2013). It is also the site of the highest solar irradiance recorded on Earth (Wierzchos et al., 2015). Moisture on the rock surface at Lomas de Tilocalar was only detected during and shortly after a rain event and the air RH is one of the lowest reported for the Atacama Desert (DiRuggiero et al., 2013; Wierzchos et al., 2013; Wierzchos et al., 2015). In contrast, the yearly average of air RH at Valle de la Luna is about twice that in Lomas de Tilocalar and the presence of a hardened layer and microrills on the calcite rock surface was indicative of dew formation (DiRuggiero et al., 2013). This is important because dew formation, together with higher RH, can significantly increase the atmospheric water budget for the calcite microbial community (Kidron, 2000; Büdel et al., 2008; DiRuggiero et al., 2013). Higher maximum temperatures in Lomas de Tilocalar could have the opposite effect by potentially increasing the rate of evapotranspiration, limiting moisture availability within the ignimbrite (Houston and Hartley, 2003).

In addition to atmospheric water availability, the structure and geochemical composition of a rock substrate are also essential determinants of its bioreceptivity (Walker and Pace, 2007; Nienow, 2009). While the mineralogy between the two substrates was significantly different, a laminated calcite layered rock and a pyroclastic rock composed of plagioclase and biotite embedded in a matrix of weakly-welded glass shards, we argue here that the major differences lied within the physical properties of the rocks (DiRuggiero et al., 2013; Wierzchos et al., 2013; Cámara et al., 2014). The calcite rocks displayed large colonized cracks and fissures, dozens of millimeters deep, all connected to the surface and providing direct access to surface water and increased light illumination. In contrast, the bottle-shaped pores found in the ignimbrite represented a limited colonization space. Not all the pores were colonized, as observed by SEM-BSE, probably because not connected to the surface. In addition, the dark varnish cover of the ignimbrite significantly decreases the incoming PAR radiation, which explains the very narrow colonization zone right underneath the rock’s surface we observed.

Taken together, these observations point to a calcite environment far less extreme than that of the ignimbrite. This is substantiated by biodiversity estimates from 16S rRNA gene sequences, and from the metagenome data, showing a higher taxonomic diversity for the calcite than for the ignimbrite community. While both rock communities were dominated by members of the *Cyanobacteria*, we found a significantly higher proportion of chlorophototrophs (>20% difference) in the ignimbrite than the calcite community. In a previous study of halite nodules, we reported a higher fraction of the cyanobacterium *Halothece* in samples collected in the driest and more extreme area of the desert than in Salars exposed to periodic fog events (Robinson et al., 2015). It was suggested that a higher proportion of photoautotrophs was required to support the associated heterotrophic populations because of lower primary productivity under water stress (Robinson et al., 2015). This might also be the case for the differences in chlorophototrophs abundance we observed between the calcite and ignimbrite rocks.

It is important to point out that the metagenome for each community was obtained with DNA extracted from four individual rocks each. The 16S rRNA gene data was obtained for each of the individual rocks and supported the taxonomic assignments of the metagenomes. Using the metagenomic data, we identified the dominant cyanobacteria in the calcite substrate as *Gleocapsa* whereas previous work assigned them to the *Chroococcidiopsis* (DiRuggiero et al., 2013). This discrepancy arose because the *Gleocapsa* and *Chroococcidiopsis* taxa are closely related, we use metagenome data, and different taxonomic assignment algorithms produces different data. It is also possible that these belong to a new genus that is neither *Gleocapsa* or *Chroococcidiopsis*.

The identification of photosynthesis and carbon fixation pathways from the metagenomes confirmed that primary production was mainly carried out via photosynthesis. However, in contrast to Antarctic cryptoendolithic communities, we did not detect any genes for chemolithoautotrophy (Wei et al., 2015). Genes for diazotrophy were also absent from both metagenomes and the major sources of nitrogen for the communities were potentially ammonia and nitrates. These later nitrogen sources might not be limiting in the Atacama Desert where high nitrate concentration from atmospheric origin (Michalski et al., 2004) have been reported. This might explain the absence of nitrogen fixation in endoliths from this desert in contrast to endolithic niches in the McMurdo Dry Valleys (Chan et al., 2013; Wei et al., 2015).

The metabolic genes and pathways annotated from the calcite and ignimbrite metagenomes revealed that the rock substrate is a stressful environment with its own challenges. Nutrient limitation was suggested by a number of stress adaptation strategies, including an abundance of protein encoding genes for two major pathways for phosphate acquisition. In cyanobacteria, and a number of other bacteria, the Pho regulon, involved in transport and metabolism of phosphonates, is controlled by environmental inorganic phosphate levels and mediates an adaptive response to phosphate starvation (Monds et al., 2006; Adams et al., 2008). Additionally, phosphonates can also be used as an alternative source of phosphorus under limiting conditions (Dyhrman et al., 2006; Adams et al., 2008). The large number of genes involved in carbon starvation found in the metagenomes is also indicative of periods of low-nutrient stress in the rock environment. These genes play a role in long-term survival under extended periods of starvation (Kolter et al., 1993), provide resistance to additional stresses such as oxidizing agent and near-UV radiation (Watson et al., 1998), and might give a competitive advantage to the cells once growth resumes (Zambrano et al., 1993).

Adaptation to desiccation in the rock environment involved a number of strategies for desiccation avoidance and protection via the biosynthesis and accumulation of organic osmolytes. The metagenome analyses revealed metabolic pathways for a broad spectrum of compatible solutes that have been found across bacterial phyla (da Costa et al., 1998; Roberts, 2005). These solutes are thought to contribute to desiccation tolerance because in high concentrations they generate low water potentials in the cytoplasm without incurring metabolic damage (Yancey, 2005; Oren, 2007). Of note, is the high abundance of genes for polyol metabolism, which in algae, fungi, plants, and a few bacteria, is a key part of a biochemical protective strategy against water loss (da Costa et al., 1998; Holzinger and Karsten, 2013). Functional categories for osmoregulation were more diverse in the calcite community but were not from the most abundant taxa, suggesting that the calcite functional diversity was driven by the taxonomic diversity of the less abundant taxa in the community. We were not able to identified genes for EPS biosynthesis in the metagenomes but microscopic observations of the endolithic communities revealed abundant EPS in the calcite but not in the ignimbrite microbial assemblages. In the calcite environment with more liquid water available, and frequent wetting-drying cycles, EPS might play a significant role in water retention, as demonstrated in similarly dry environments (Hu et al., 2012). In contrast, in the ignimbrite, where the unique source of liquid water are very scarce rain events and complete dehydration periods lasting as long as 9 months, the environmental conditions might too harsh for EPS effectiveness and for the energy expenditure required for their biosynthesis.

Cyanobacteria are known to produce a wide range of secondary metabolites including PKS, NRPS, and antibiotics (Leao et al., 2012; Puglisi et al., 2014). One of the major differences, at the functional level, between the calcite and ignimbrite metagenomes was for secondary metabolites pathways. We found a considerably higher number of NRPs and PKS related genes in the ignimbrite metagenome, and the draft genome of some of the most abundant cyanobacteria from the ignimbrite community, than in the calcite community. Biosynthetic pathways for other secondary metabolites included those for alkaloids, involved in host fitness and protection (Leflaive and Ten-Hage, 2007), phenylpropanoids, which have been shown to enhance antioxidant activity and tolerance to stress in cyanobacteria (Singh et al., 2014), phenazines known to contribute to behavior and ecological fitness (Pierson III and Pierson, 2010), and phytoalexins with antimicrobial and often antioxidative activities (Patterson and Bolis, 1997). The synthesis of these secondary metabolites is often directed toward protection from radiation, allelopathy, resource competition, and signaling (Leao et al., 2012; Puglisi et al., 2014) and, in the case of the ignimbrite, might be an indication of the fierce competition for scarce resources and limited colonization space in the small pores of this substrate rock. Indeed the production of small antimicrobial compounds has been reported as a strategy for eliminating prior residents before colonization of a new space by a number of microorganisms (Hibbing et al., 2010). Additionally, SEM-BSE images of colonized pores from the ignimbrite rock revealed cell decayed cells in some of the pores, potentially reflecting the effects of antimicrobial compounds or resource competition. In contrast the chasmoendolithic environment of the calcite provided more physical space and therefore may promote interactions between members of the community for metabolic cooperation. In addition to antimicrobials, database searches revealed that the closest chemical structures for some of the NRPs and PKS gene clusters found in both communities were for siderophores. Iron is an essential element for most organisms because of its essential role in redox enzymes, membrane-bound electron transport chains, but also in photosynthesis (Andrews et al., 2003). The abundance of siderophores in the rock communities, and the enrichments in genes for iron acquisition in the cyanobacteria draft genome when compared to the reference genome from *N. punctiforme*, strongly suggest iron starvation. This was emphasized in the ignimbrite community with a large number of sequence reads assigned to gene clusters for siderophores and a ferrichrome-iron receptor. Furthermore, the mycosporine gene cluster found in the calcite cyanobacteria, but not the ignimbrite, may indicate differences in UV radiation stress experienced by the two communities (Garcia-Pichel et al., 1993; Oren, 1997).

Environmental stress is a major driver of microbial diversity and this is reflected in the functional annotation of metagenomes from Atacama Desert endoliths. Our analysis is reflective of adaptations to desiccation, osmotic stress, low nutrients, iron deficiency, and in the case of the ignimbrite, of a fierce battle for colonization space and resources among community members. The relative abundance of phototrophs in each community is likely indicative of the *in situ* level of primary production, a hypothesis that might be tested by careful field measurements of microbial activity.

## Acknowledgements

This work was funded by grant EX0B08–0033 from NASA and grant NSF-0918907 from the National Science foundation to JDR and by MINECO (Spain) grants CGL2013–42509P to JW, CA, JDR and OA.

---

| Compound | Biosynthetic gene cluster<br>(antiSMASH) | Family<br>(NORINE) | Activity<br>(NORINE) |
| --- | --- | --- | --- |
| a | NRPS | ACV | antibiotic |
| b | NRPS-T1PKS Microcystin<br>(siderophore) | coronatine | toxin |
| c | NRPS-T1PKS<br>Anabaenopeptin | largamide | protease<br>inhibitor |
| d | NRPS Xenoamicins |  |  |
| e | NRPS | pyoverdin | siderophore |
| f | NRPS-T1PKS | pyoverdin | siderophore |
| g | NRPS Nostophycin | coronatine | toxin |
| i | NRPS similar to bacteriocins |  |  |
| I | NRPS siderophore | coronatine | toxin |

